# Chronic Social Isolation Reshapes the Metabolome in *Drosophila*

**DOI:** 10.64898/2026.09.24.754212

**Authors:** Doris Migliaccio, Carmen Spicer, Andrew D. Nguyen, Aabha Vora, Zhaoyue Zhang, Wanhe Li

## Abstract

Sleep, feeding, and metabolism are tightly coordinated outputs of the circadian system. Previous studies showed that chronic social isolation reduces sleep and increases feeding in *Drosophila*. Here, untargeted metabolomic profiling revealed that chronic social isolation induces a distinct metabolomic signature spanning amino acids, carbohydrates, and lipids, establishing a foundation for understanding the physiological consequences of social isolation.

## Introduction

Social isolation is increasingly recognized as a major determinant of health, particularly following the COVID-19 pandemic^1,2^. Large-scale epidemiological studies have identified loneliness as a significant risk factor for morbidity and mortality, with effect sizes comparable to established risk factors such as obesity and smoking. Human studies have also linked social isolation to metabolic disorders, including obesity and type 2 diabetes^3-5^. Animal studies have demonstrated that social isolation alters gene expression in metabolism pathways and affects metabolic processes across multiple tissues^6-11^. However, direct characterization of how social isolation reshapes metabolism at a systems level remains limited.

Sleep, feeding, and metabolism are tightly coordinated outputs of the circadian system^12,13^. In *Drosophila melanogaster*, chronic social isolation induces some of the most robust behavioral changes^9,14-21^, including reduced sleep and increased food consumption, making it a powerful model to investigate how social experience influences internal physiological state^9^. Previous work demonstrated that chronic social isolation leads to reduced sleep and increased food consumption despite uninterrupted access to nutrients, together with transcriptional changes consistent with altered metabolic regulation^8,9,22^. These findings suggest that social isolation may broadly influence metabolic physiology.

Metabolomic profiling has become a powerful systems-level approach for investigating metabolism and circadian physiology^23-26^. However, whether chronic social isolation produces corresponding changes in the metabolome remains unknown. Here, we performed untargeted metabolomic profiling of chronically group-housed and socially isolated *Drosophila*. We show that chronic social isolation produces a distinct metabolomic signature characterized by coordinated changes in amino acids, central carbon metabolites, and lipids. These metabolic changes are reproducible across trials and genetic backgrounds and further distinguish intermediate social conditions, establishing metabolomics as a framework for investigating the physiological consequences of social isolation.

## Results

To determine whether chronic social isolation affects systemic metabolism, we performed untargeted metabolomic profiling using the wild-type isogenic strains 2202u, maintained under group-housed or isolated conditions for 7 days^9^. Chronic social isolation produced coordinated changes across multiple metabolite classes, including amino acids, central carbon metabolites, and lipids (Fig. 1 and Supplementary Fig. 1).

**Figure 1.**
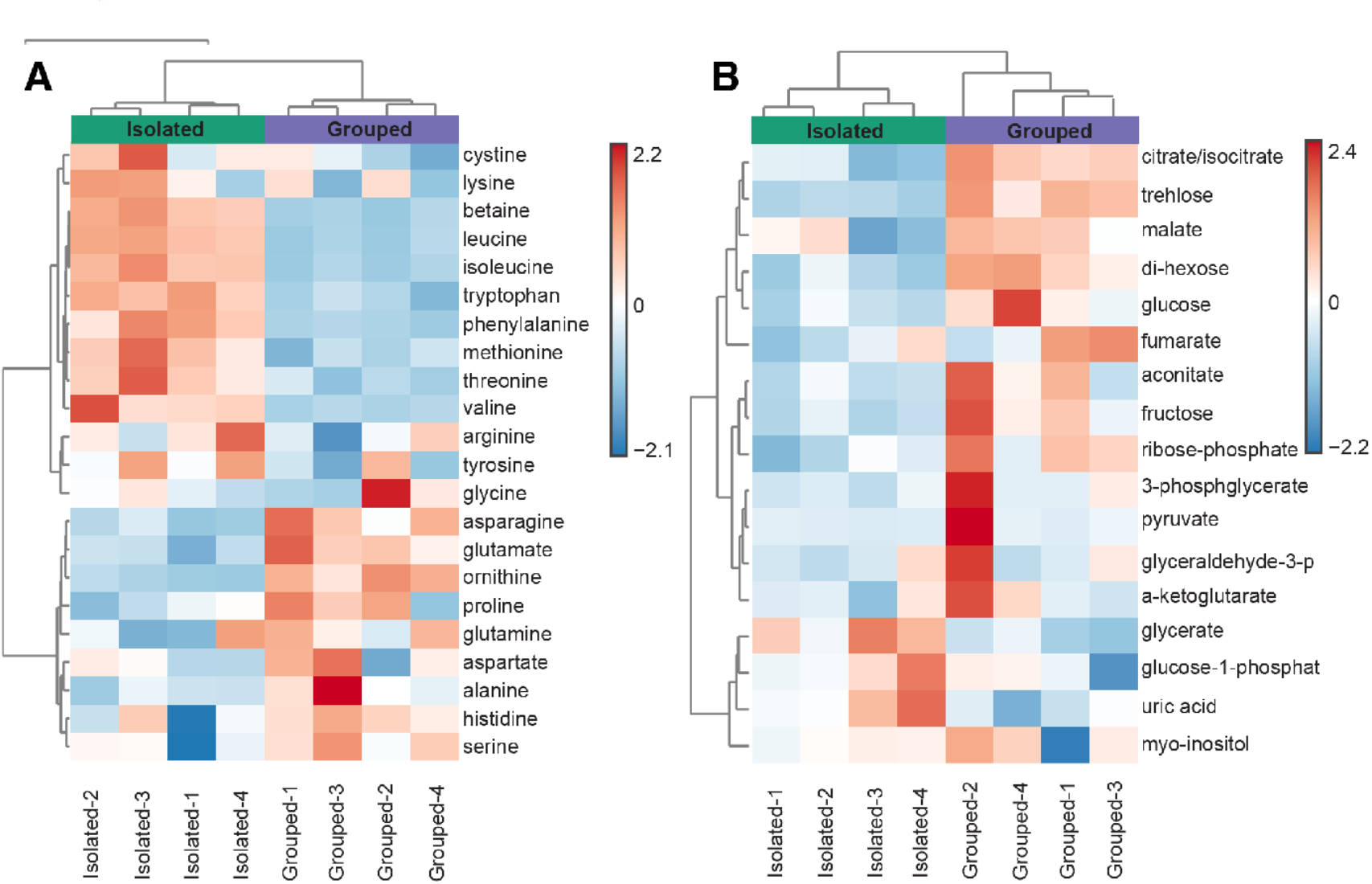
Chronic social isolation induces widespread metabolomic changes in 2202u flies. (**A)** Heatmap showing unsupervised hierarchical clustering of amino acid metabolites from 2202u flies after 7 days group housing or social isolation. **(B)** Heatmap showing unsupervised hierarchical clustering of central carbon metabolites from 2202u flies after 7 days group housing of social isolation. Columns represent individual biological replicates and rows represent metabolites. Metabolite abundance is displayed as row-scaled Z-scores.

Analysis of amino acid profiles revealed a divergence between essential and non-essential amino acids (Fig. 1A). Several essential amino acids, including the branched-chain amino acids isoleucine and leucine, as well as methionine, phenylalanine, threonine, and tryptophan, were elevated in isolated flies. In addition, betaine was also increased. In contrast, non-essential amino acids including glutamate, asparagine, and ornithine were reduced. These findings indicate that chronic social isolation is associated with a redistribution of amino acid pools.

We next examined metabolites associated with central carbohydrate metabolic pathways (Fig. 1B). Isolated flies exhibited reduced levels of several key metabolites, including trehalose, the principal circulating sugar in *Drosophila*, as well as di-hexose species. In addition, glycerate, a glycolytic intermediate, and citrate/isocitrate, key intermediates of the tricarboxylic acid cycle, were decreased. These findings indicate that chronic social isolation is associated with reduced levels of circulating sugars and central carbon metabolites.

To assess how lipid metabolism is affected by social isolation, we next performed untargeted lipidomic profiling. Chronic social isolation resulted in widespread changes in lipid abundance. Numerous lipid species, including those of phosphatidylinositol, phosphatidylethanolamine, phosphatidylcholine, phosphatidylglycerol, and ceramide were reduced in isolated flies, whereas only a limited subset of lipid species increased (Supplementary Fig. 1). Importantly, heatmaps of amino acid, central carbon, and lipid metabolites showed that unsupervised hierarchical clustering clearly separated group-housed and socially isolated flies, with biological replicates clustering tightly within each condition, demonstrating the robustness and reproducibility of the metabolomic differences. These findings demonstrate that chronic social isolation is associated with coordinated changes across amino acid, carbohydrate, and lipid metabolism, producing a distinct metabolomic signature.

To determine whether the metabolomic changes observed in the 2202u strain were reproducible and conserved across genetic backgrounds, we performed a second metabolomic analysis using an independent cohort of 2202u flies together with another commonly used wild-type strain, Canton-S (CS), both maintained under group-housed or isolated conditions for 7 days (Fig. 2 and Supplementary Fig. 2). Unsupervised hierarchical clustering clearly reproduced the separation between group-housed and socially isolated 2202u flies. In the Canton-S strain, clustering also generally separated the two housing conditions, although one isolated biological replicate clustered with the group-housed samples. Consistent with this pattern, the magnitude of metabolite changes was generally smaller in Canton-S than in 2202u flies. The clustering further revealed a clear separation between the 2202u and Canton-S strains, indicating that genetic background contributes substantially to variation in the metabolome (Fig. 2A). Nevertheless, many of the changes in amino acid and central carbon metabolism were also observed in Canton-S flies (Fig. 2B, C), indicating that the metabolomic response to chronic social isolation is not unique to the 2202u genetic background. Together, these findings demonstrate that the metabolomic response to chronic social isolation is reproducible, conserved across genetic backgrounds, and exhibits quantitative strain-dependent differences. In contrast to the polar metabolite profiles, lipidomic clustering was largely driven by genetic background, with 2202u and Canton-S samples forming two clearly separated clusters regardless of housing condition (Supplementary Fig. 2).

**Figure 2.**
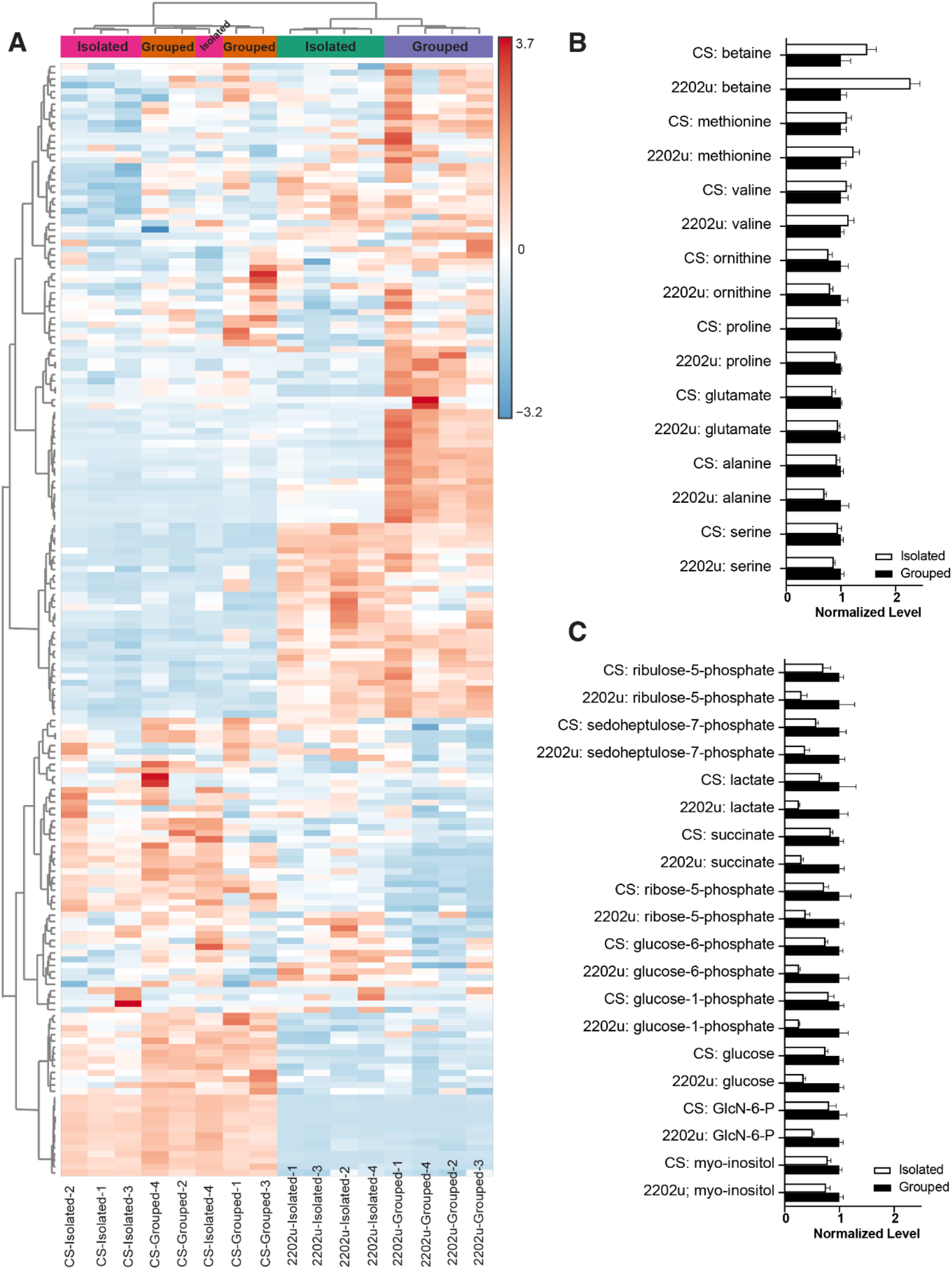
Metabolomic responses to chronic social isolation across genetic backgrounds. **(A)** Heatmap showing unsupervised hierarchical clustering of polar metabolites from 2202u and Canton-S (CS) flies after 7 days of group housing or social isolation. Columns represent individual biological replicates and rows represent metabolites. Metabolite abundance is displayed as row-scaled Z-scores. **(B)** Relative abundances of selected amino acid metabolites in group-housed and socially isolated 2202u and CS flies. **(C)** Relative abundances of selected central carbon metabolites in group-housed and socially isolated 2202u and CS flies. Metabolite levels are normalized to the mean value of the corresponding group-housed samples within each strain, which was set to 1.

To determine whether metabolomic changes reflect graded differences in social environment, we included an intermediate Double housing condition, in which two flies were housed together, and compared metabolite profiles among Grouped, Double, and Isolated 2202u flies (Fig. 3). Examination of individual metabolites identified distinct purine nucleotide signatures that were associated with each housing condition. Among these, xanthosine nucleotides exhibited a striking pattern, with Grouped flies showing the highest levels of xanthosine monophosphate (XMP), Isolated flies exhibiting the highest levels of xanthosine diphosphate (XDP), and Double flies showing the highest levels of xanthosine triphosphate (XTP). Together, these findings demonstrate that metabolomic profiling distinguishes not only Group and Isolated flies but also the intermediate Double condition, indicating that distinct social environments are associated with unique metabolomic states.

**Figure 3.**
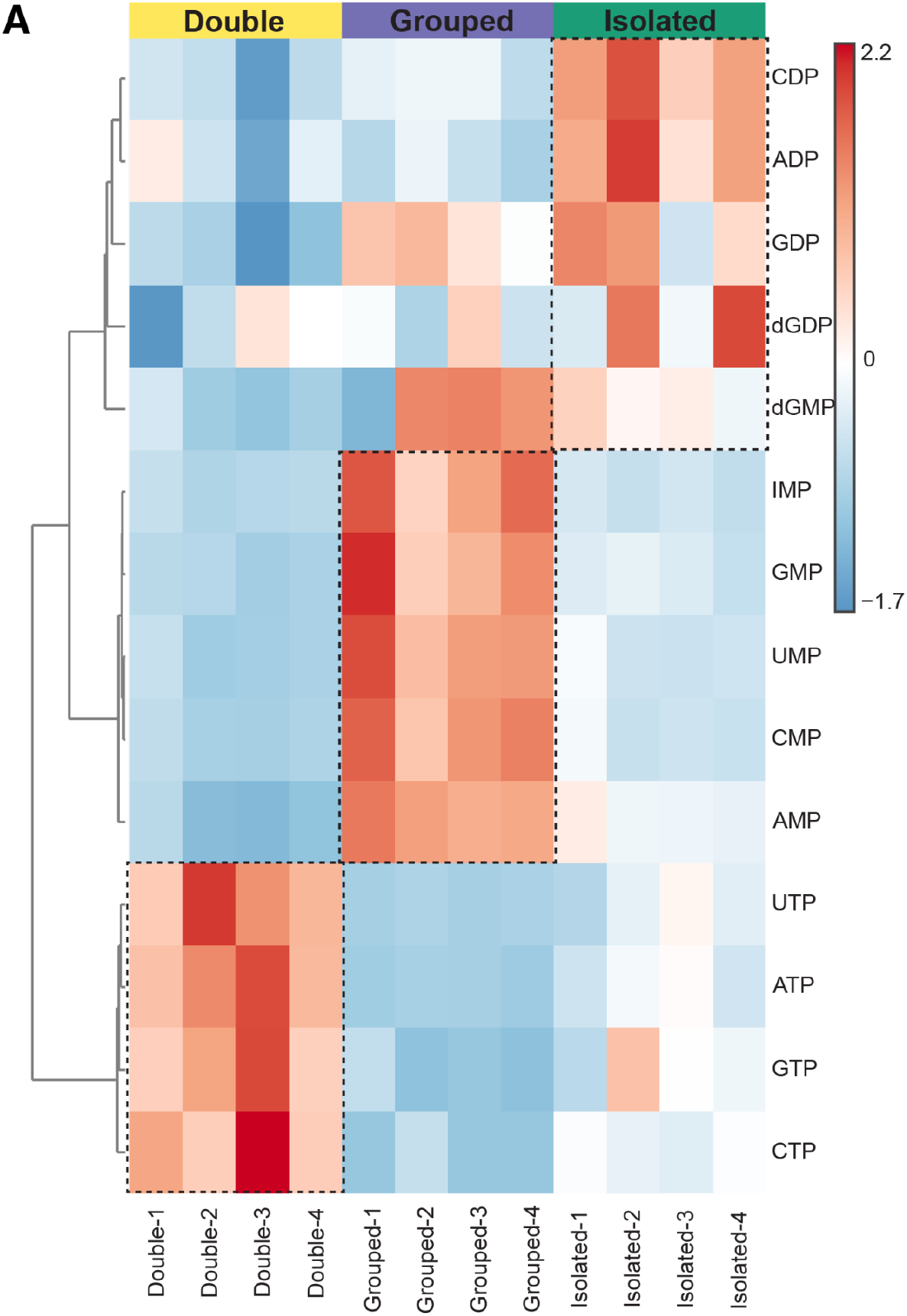
Distinct xanthosine nucleotide signatures associated with different social environments. Heatmap showing unsupervised hierarchical clustering of xanthosine monophosphate (XMP), xanthosine diphosphate (XDP), and xanthosine triphosphate (XTP) in Grouped, Double, and Isolated 2202u flies after 7 days of housing. Columns represent individual biological replicates and rows represent metabolites. Metabolite abundance is displayed as row-scaled Z-scores.

## Discussion

Chronic social isolation has been extensively studied through behavioral, physiological, and transcriptomic approaches. Here, we show that it is also associated with widespread metabolomic remodeling. Across independent experiments, chronic social isolation produced coordinated changes in amino acids, central carbon metabolites, and lipids. These metabolic alterations were reproducible in an independent cohort of 2202u flies and were also observed in the widely used Canton-S strain, although with smaller effect sizes. Furthermore, introducing an intermediate social condition demonstrated that distinct social environments are associated with distinct metabolomic states.

An important feature of this study is that these metabolomic changes were induced by a remarkably simple environmental manipulation. Flies were maintained under identical dietary and environmental conditions, differing only in the degree of social interaction.

The widespread metabolic differences observed under these conditions highlight the profound influence of social experience on systemic physiology. These findings are also consistent with our previous demonstration that chronic social isolation induces sleep loss and hyperphagia, defining a distinct brain state that resembles persistent hunger despite uninterrupted nutrient availability^9^.

The physiological consequences of social isolation have traditionally been characterized through behavioral phenotypes, gene expression profiling, and neural circuit analyses. Our findings suggest that metabolomic profiling provides a complementary systems-level approach for investigating how social experience influences organismal physiology. At the same time, the quantitative differences observed between 2202u and Canton-S strains indicate that the magnitude of the metabolomic response is influenced by genetic background.

Together with its powerful genetics, well-defined behavioral paradigms, and capacity for high-throughput experimentation, *Drosophila melanogaster* provides an efficient model for integrating metabolomics with behavioral and molecular analyses. This framework should facilitate future studies investigating how social experience is translated into physiological and metabolic states.

## Methods

### *Drosophila* Strains and Culture

*Drosophila melanogaster* stocks were raised on standard medium (cornmeal/yeast/molasses/agar) at 21.5°C under 12-h light: 12-h dark (LD) cycles. The wild-type isogenic strains 2202u, an isogenic derivative of Canton-S carrying the *w*^*1118*^(*isoCJ1*) allele^27^, and Canton-S (CS) were used. All experiments were conducted in male animals.

### Group Enrichment/Social Isolation

Newly eclosed flies were collected and kept in standard fly food bottles (∼200 flies per bottle) for 3-5 days to acquire social experience. Male flies were then sorted into standard fly food vials: 1 fly per vial for social isolation (Isolated) and 20 flies per vial for group enrichment (Grouped). For all experiments, the fly sorting day was considered day 0. Social isolation was conducted in a chronic manner, with flies being isolated for a total of 7 days. Group flies were parallelly exposed to group enrichment for 7 days.

### Sample Collection for Metabolomics

Following 7 days of group enrichment or social isolation, flies were collected for metabolomic analysis during ZT1-ZT2. For each condition, 20 flies per sample were pooled, with four independent biological replicates per condition. Samples were rapidly collected using liquid nitrogen and processed to minimize metabolic perturbation.

### Polar Metabolites Extraction

Intracellular metabolites were extracted using a cold methanol-based extraction protocol. Briefly, pooled fly samples were homogenized in 800 μl of 80% (v/v) aqueous methanol using ceramic beads. Samples were vortexed and incubated at 4 °C on a rotator for 1 hour, followed by centrifugation at 20,000g for 15 minutes. The resulting supernatant containing polar metabolites was collected for downstream analysis.

### Non-polar Metabolites Extraction

For lipidomic (non-polar metabolites) analysis, metabolites were extracted using a modified Folch two-phase extraction method. Tissue samples were homogenized in cold chloroform and methanol, followed by phase separation with the addition of water. This procedure generated a biphasic system consisting of a methanol phase containing polar metabolites and a chloroform phase containing non-polar metabolites. Phases were carefully separated, avoiding the interphase, and dried prior to analysis. All extraction steps were performed on ice or at 4 °C to preserve metabolite integrity. Extracted samples were dried using a centrifugal evaporator or nitrogen stream and stored at −80 °C until LC-MS analysis.

### Liquid Chromatography-Mass Spectrometry (LC-MS) Analysis

Polar metabolites were analyzed using liquid chromatography–mass spectrometry (LC-MS). Separation was performed on a ZIC-pHILIC column (150 × 2.1 mm, 5 μm particle size) coupled to a Thermo Vanquish UPLC system and a Q Exactive Orbitrap mass spectrometer equipped with a heated electrospray ionization (HESI) source. Mobile phase A consisted of 20 mM ammonium carbonate with 0.1% ammonium hydroxide (pH 9.3), and mobile phase B consisted of acetonitrile. Metabolites were separated using a gradient from 90% to 40% B over 22 minutes, followed by equilibration to initial conditions. The flow rate was maintained at 0.15 ml min^−1^. Mass spectrometry was performed in polarity-switching mode with the following parameters: spray voltage 3.0 kV, capillary temperature 275 °C, source temperature 250 °C, sheath gas flow 40 arbitrary units, and auxiliary gas flow 15 arbitrary units. Full MS scans were acquired at 70,000 resolution with an automatic gain control (AGC) target of 1 × 10^6^, a maximum injection time of 80 ms, and a scan range of m/z 55–825.

### Lipidomic Analysis

Non-polar metabolites were analyzed using untargeted LC-MS/MS lipidomics. Lipid species were separated using reverse-phase chromatography (C18) and detected in both positive and negative ionization modes. Raw data were processed and annotated using lipid identification software LipidSearch, enabling detection of a broad range of lipid classes including phospholipids, sphingolipids, and fatty acids.

### Data processing and analysis

Metabolomic data were processed using MetaboAnalyst for normalization, statistical analysis, and pathway-level interpretation. All experiments were performed with four independent biological replicates.

## Data Availability

All data supporting the findings of this study are available within the paper and its Supplementary Information.

## Acknowledgements

We thank Michael Young for sharing *Drosophila* strains and equipment. We thank members of the Li laboratory for discussions. Metabolomic data was generated by the Proteomics Resource Center at The Rockefeller University (RRID:SCR_017797) using instrumentation funded by the Sohn Conferences Foundation and the Leona M. and Harry B. Helmsley Charitable Trust. This work was supported by the Cancer Prevention and Research Institute of Texas [RR220021] and the National Institute of General Medical Science [GM150832] to W. L.. Z. Z was supported by the Helen Hay Whitney Fellowship.

## Author Contributions

Conceptualization: D.M., Z.Z. and W.L.; sample collection and data curation: C.S., A.N., A.V., Z.Z. and W.L.; formal analysis: Z.Z.; funding acquisition: W.L.; supervision: W.L.; writing-draft: D.M. and W.L.; writing-editing: D.M., Z.Z., and W.L..

## Competing Interests

The authors declared no potential conflicts of interest with respect to the research, authorship, and/or publication of this article.

## Supplementary Materials

**Supplementary Figure 1.**
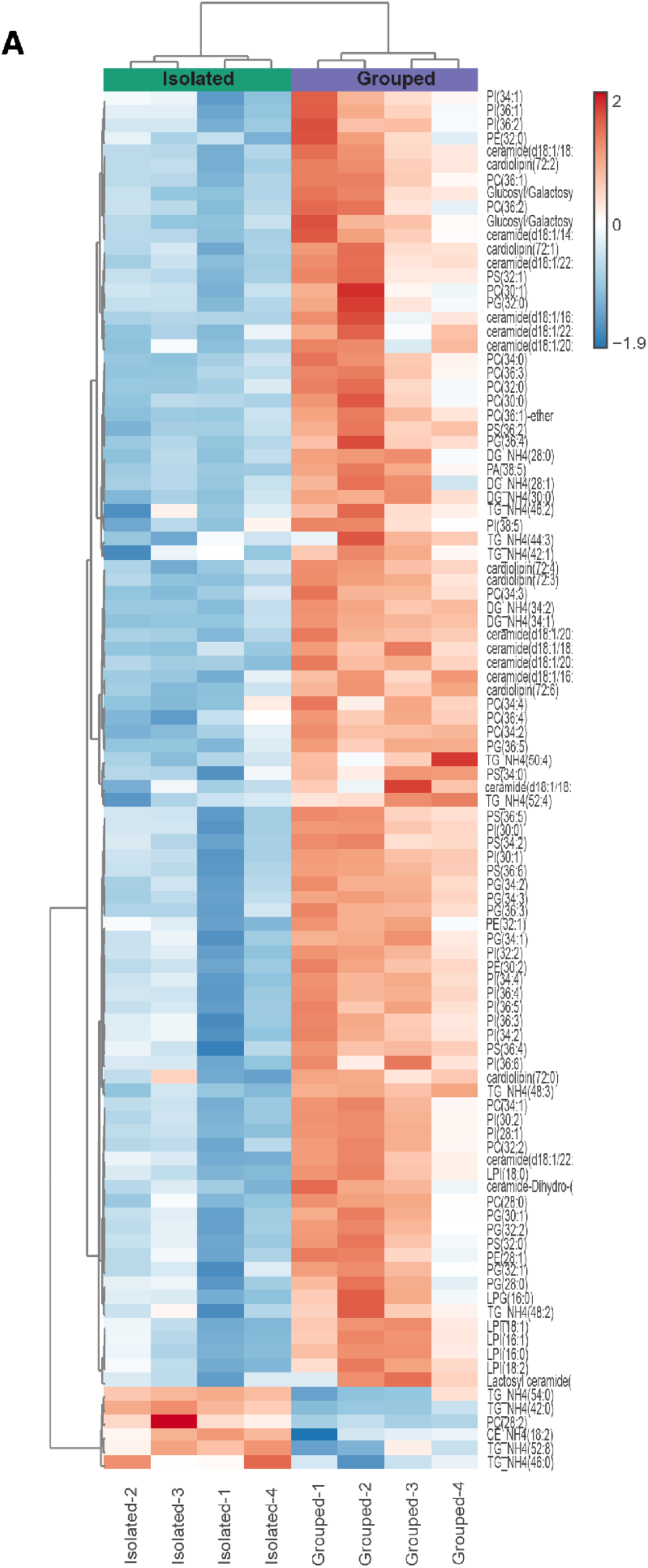
Hierarchical clustering of the top 100 significantly altered lipids following chronic social isolation. Heatmap showing unsupervised hierarchical clustering of the 100 most significantly altered lipid species between group-housed and socially isolated 2202u flies after 7 days of housing. Columns represent individual biological replicates and rows represent lipid species. Lipid abundance is displayed as row-scaled Z-scores.

**Supplementary Figure 2.**
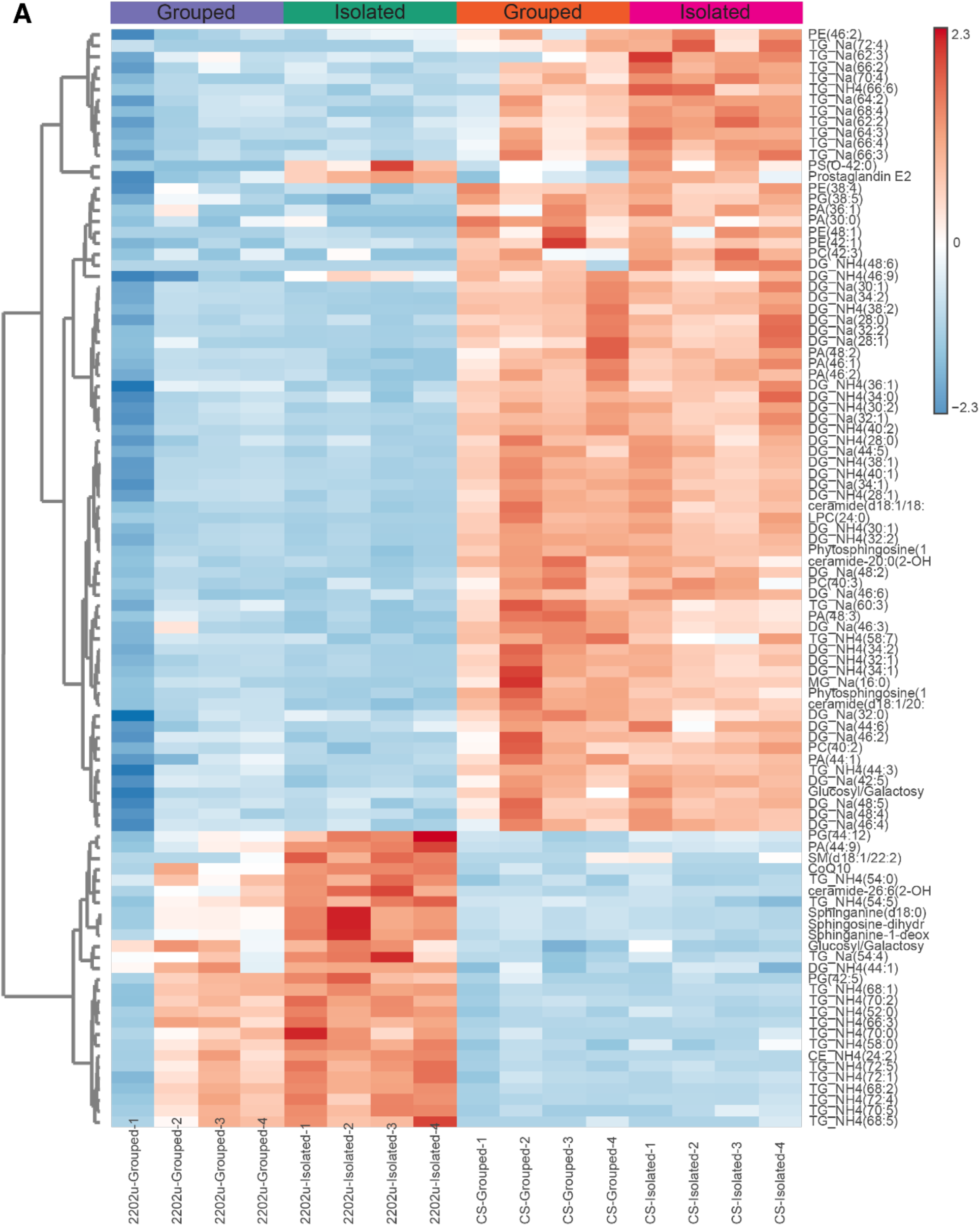
Hierarchical clustering reveals distinct lipidomic profiles in 2202u and Canton-S flies. Heatmap showing unsupervised hierarchical clustering of the top 100 differentially abundant lipid species from group-housed and socially isolated 2202u and Canton-S (CS) flies after 7 days of housing. Columns represent individual biological replicates and rows represent lipid species. Lipid abundance is displayed as row-scaled Z-scores.

## Notes

### Competing Interest Statement

The authors have declared no competing interest.

